# Hedgehog pathway members Patched and Costal-2 exhibit differences in overgrowth autonomy in *Drosophila melanogaster*

**DOI:** 10.1101/2021.04.09.439246

**Authors:** Shannon L. Moore, Frank C. Adamini, Erik S. Coopes, Dustin Godoy, Shyra J. Northington, Jordan M. Stewart, Richard L. Tillet, Kayla L. Bieser, Jacob D. Kagey

## Abstract

Genetic screens are used in *Drosophila melanogaster* to identify genes key in the regulation of organismal development and growth. These screens have defined signaling pathways necessary for tissue and organismal development which are evolutionarily conserved across species, including *Drosophila*. Here we have used a Flp/FRT mosaic system to screen for conditional regulators of cell growth and cell division in the *Drosophila* eye. The conditional nature of this screen utilizes a block in the apoptotic pathway to prohibit the mosaic mutant cells from dying via apoptosis. From this screen, we identified two different mutants that mapped to the Hedgehog signaling pathway. Previously, we described a novel *Ptc* mutation and here we add to the understanding of disrupting the Hh pathway with a novel allele of *Cos2*. Both of these Hh components are negative regulators of the pathway, yet they depict mutant differences in the type of overgrowth. *Ptc* mutations lead to overgrowth consisting of almost entirely wild type issue (non-autonomous overgrowth), while the *Cos2* mutation results in tissue that is overgrown in both the mutant and wild type clones (both autonomous and non-autonomous). These differences in tissue overgrowth are consistent in the *Drosophila* eye and wing. The observed difference is correlated with a different pattern of deregulation of Mad, the downstream effector of DPP signaling. This finding provides insight into pathway specific differences that may help to better understand intricacies of developmental processes and human disease.

## Introduction

*Drosophila melanogaster* is a long-established model system with particular usefulness in understanding the genetic mechanisms underlying many life processes, including the genetic regulation of growth and development. A large proportion of the genes that regulate growth in developing tissues across species, including humans, were first identified using genetic screens in *Drosophila* (Stone et al., 1996). The Flp/FRT genetic system has been used in genetic screens to identify genes that are involved in regulating growth and differentiation by analyzing the ratios of mutant to wild type tissue within the *Drosophila* eye. Genes identified in these screens are frequently implicated in human diseases, and a number of these genes were initially discovered in *Drosophila* and later found to be disease causing in human (Hariharan and Bilder, 2006; Harvey et al., 2003; Moberg et al., 2001; Wu et al., 2003).

In developing our genetic screen, we hypothesized that a subset of mutations would lead to the removal of mutant cells through apoptosis (when homozygous), thus causing specific mutantations to be missed in the initial iterations of the Flp/FRT screen (Golic and Lindquist, 1989; St Johnston, 2002). Since the inhibition of apoptosis remains a key step during carcinogenesis for cancer cells to avoid a primary defense mechanism (Hanahan and Weinberg, 2000), this inhibition remains an important genetic process to understand from a disease standpoint. Therefore, we conducted a second-generation conditional Flp/FRT EMS screen on chromosome 2R in a genetic background blocking apoptosis, to identify conditional regulators of cell growth and division (Kagey et al., 2012). A number of growth mutants conditional on a block in apoptosis have been previously identified, including *scribbled*, providing further rationale for the screen (Brumby and Richardson, 2003).

From this screen we have previously reported on a number conditional regulators of cell growth and cell division including *CPA* and *Shn*, demonstrating the occurrence of these conditional growth mutants (Bieser et al., 2019; Cosenza and Kagey, 2016). Here we describe the nature of the over-growth caused by two of these mutants identified in the screen that are both part of the Hedgehog signaling pathway, *Patched (Ptc)* and *Costal-2 (Cos2)*.

The Hedgehog pathway (Hh) is a highly conserved signaling pathway essential to organismal development (Bellusci et al., 1997; Goodrich et al., 1996). This pathway was first identified for its role in segment polarity during *Drosophila* embryo development (Nusslein-Volhard and Wieschaus, 1980). Since that discovery, Hh has been found to be a critical pathway in regulating cell differentiation and growth across model organisms. In humans, mutations in the Hh pathway have been linked to both developmental abnormalities and disease later in life, making it a relevant diagnostic and therapeutic target (Hahn et al., 1996).

The Hh pathway starts with the Hedgehog (Hh) ligand binding to the extracellular receptor Patched (Ptc). When unbound, Ptc inhibits Smoothened (Smo). The binding of Hh stops suppression of Smoothened, leading to downstream activation of proteins including Protein Kinase A (PKA) and the kinesin Costal-2 (Cos2) (Jiang and Hui, 2008). When activated, PKA/Cos2 cleave the transcriptional activator Ci, allowing its translocation into the nucleus, where it drives gene transcription (Wang and Holmgren, 2000). Ci has several known transcriptional targets including *Ptc*, which then serves as a negative feedback loop for the pathway (Xiong et al., 2015).

Previously, we reported on one mutant from the screen, *Ptc*^*B.2.13*^, that serves as a conditional regulator of overgrowth (Kagey et al., 2012). The *Ptc*^*B.2.13*^ mutation to leads to non-autonmous overgrowth driven by the non-autonomous activity of Yorkie and Mad in cells that are adjacent to the *Ptc*^*B.2.13*^ clones. Complementing these findings was a companion study utilizing the *GMR-Hid* mosaic screen conducted by the Bergmann lab (Fan and Bergmann, 2008).

Here, we add to the knowledge of the interplay between growth regulation, tissue development, and cell survival in the Hh pathway. We find that Cos2 is also a Hh pathway conditional regulator of cell growth, dependent upon a block in the canonical apoptosis pathway. However, despite both Ptc and Cos2 serving as negative regulators of the Hh signaling pathway, we find differences in the autonomous nature of the overgrowth: *Ptc* mutant clones drive a non-autonomous overgrowth while *Cos2* mutant clones lead to both an autonomous and non-autonomous overgrowth. These findings provide insight into how disruptions to different points of the Hh signaling pathway lead to different consequences for growth and development.

## Methods

### Conditional Genetic Screen for Regulators of Cell Growth and Division

An EMS screen was conducted on chromosome 2R utilizing Flp/FRT system in the *Drosophila* eye. This screen utilized *FRT42D, Dark*^*82*^ as a starting chromosome to identify mutations that disrupted cell division and growth, dependent on the *Dark*^*82*^ block of apoptosis (Kagey et al., 2012). The *Dark*^*82*^ allele utilized has been shown to block developmental apoptosis when homozygous (Akdemir et al., 2006). From the screen 137 mutants were identified and stable stocks were created. Selected growth mutants were mapped via complementation mapping to the Bloomington 2R Deficiency kit (Cook et al., 2012). Previously we mapped a novel allele of *Patched (Ptc*^*B.2.13*^*) (Kagey et al*., *2012)*. From the same screen here we report a novel allele of *Costal-2* (*Cos2*^*F.1.4*^). The mutant stock *Cos2*^*F.1.4*^ failed to complement two previously characterized alleles of *Cos2, Cos2*^*k16101*^ and *Cos2*^*H29*^ (Christiansen et al., 2013; Roch et al., 1998).

To identify the specific molecular lesion causing driving the growth phenotypes in *Ptc* and *Cos2* PCR primers were designed for the exons of each gene. DNA was isolated utilizing Li/Cl, KAc from heterozygous *Ptc*^*B.2.13*^ and *Cos2*^*F.1*.4^ animals. Following successful PCR amplification, samples were sent for Sanger sequencing to identify heterozygous point mutations in the chromatograms (Genewiz, New Jersey).

### Genetics

In addition to the *Dark*^*82*^ allele (Akdemir et al., 2006), the following genotypes were used in these experiments *Ey-Flp;FRT42D* (BDSC), *Ey-Flp; FRT42D, ubi-GFP, Ey-Flp; FRT42D, M(2)* (BDSC), *UBX-Flp; FRT42D, ubi-GFP* (BDSC), *Cos2*^*H29*^ allele (Christiansen et al., 2013), and Cos2^K161010^ (BDSC) (Roch et al., 1998). The 2R deficiency kit from the Bloomington *Drosophila* Stock Center was used for genetic mapping via complementation tests (Cook et al., 2012). All crosses were conducted at 25°C.

### Adult eye and wing visualization

Images were taken of the adult mosaic eyes at the same magnification (40x) under 70% ethanol to compare size of eye and ratio of mutant/wild type tissue. In cases where the cross was setup to a stock containing the pigmented *Dark*^*82*^ allele, the ‘Flp’ stock contained an unpigmented *FRT42D* chromosome. When the *Dark*^*82*^ allele was not present the ‘Flp’ stock utilized an *FRT42D* chromosome with pigmentation (*FRT42D, ubi-GFP*). Adult wings were mounted to slides in vegetable oil and imaged at the same magnification (70x) to allow comparison of size differences between genotypes. Quantification of adult wing size was done measuring pixels on Photoshop of at least 10 wings per genotype. For both wings and eyes the images were taken on an AM Scope camera (AM scope 550 MA).

### Immunohistochemistry

Imaginal eye and wing discs were dissected from wandering L3 larvae as previously described (Kagey et al., 2012; Pellock et al., 2007). Briefly, imaginal discs were fixed in 4% paraformaldehyde and permeabilized in 0.3% PBST prior to staining with antibodies.

Antibodies from Developmental Studies Hybridoma Bank: anti-Ptc (mouse, 1:40), anti-Elav (rat 1:800), and anti-Ci (rat, 1:100) were used to stain eye and wing discs. Other antibodies used were anti-phospho-Smad1/5 (pMad) (rabbit 1:100, Cell Signaling), anti-DIAP1 (mouse, 1:50) (Yoo et al., 2002), and anti-GFP (chicken, 1:100, Aves Labs). Imaginal eye and wing discs were visualized on a compound fluorescent microscope or confocal microscope using the Zen microscopy software (Zeiss microscopy).

### Analysis of fluorescent images

Imaginal eye and wing disc size and autonomy were calculated using Photoshop to measure the number of pixels in each genotype. At least 10 imaginal discs were quantified for each genotype. Autonomy was determined by calculating the percentage of non-GFP (mutant) tissue divided by the whole tissue size using pixels in Photoshop. T-tests were used to analyze the differences in autonomy between *Ptc*^*B.2.13*^ and *Cos2*^*F.1.4*^ mosaic tissue, and p values were calculated to determine the significance. For these comparisons, a p value < 0.05 was used for significance. To measure levels of pMad expression across the clonal boundaries florescent intensity was measured using the Zen microscopy line scan to measure signal intensity (Zeiss microscopy).

## Results

### Ptc and Cos2 alleles were isolated from a conditional Flp/FRT eye screen and map to genes in the Hedgehog pathway

In the genetic background of blocked apoptosis, mutants that disrupted cell growth and cell division in the mosaic eye were isolated in an EMS Flp/FRT genetic screen (Kagey et al., 2012). Previously, *Ptc*^*B.2.13*^ was mapped as a novel allele of *Ptc* via complementation mapping which resulted in a failure to complement the *Ptc*^*S2*^ allele (Ingham et al., 1991; Kagey et al., 2012).

Here we mapped the *F.1.4* mutant using homozygous lethality and complementation tests to the 2R Df kit (Cook et al., 2012). A region of failure to complement was identified in the overlapping region between the deficiencies *Df(2R)ED1715* and *Df(2R)1673* from 2R:7,326,951..7,533,553. The *F.1.4* mutant was mated to individual alleles within this region and failed to complement two previously characterized *Cos2* alleles, *Cos2*^*k16101*^ and *Cos2*^*H29*^, indicating that *Cos*^*F.1.4*^ is a novel allele of *Cos2* (Christiansen et al., 2013; Roch et al., 1998).

To further confirm the location of these mutations and to understand the molecular alterations, we conducted Sanger sequencing of both *Ptc*^*B.2.13*^ and *Cos2*^*F.1.4*^ flies. For *Ptc*^*B.2.13*^ and *Cos2*^*F.1.4*^, a double-peak in each chromatogram causing missense mutations in an amino acid conserved between flies and humans. In *Ptc*^*B.2.13*^, a T-A at 2R:8,660,339 was identified resulting in a Trp-173-Arg missense mutation (Supplemental Figure 1). In *Cos2*^*F.1.4*^, a T-A at 2R:8,660,339 was identified resulting in a Leu-951-Gln missense mutation (Supplemental Figure 1). The identified missense mutations along with the failure to complement data from established *Ptc* and *Cos* alleles establish *Ptc*^*B.2.13*^ and *Cos2*^*F.1.4*^ as mutant alleles of the Hh pathway.

### Ptc and Cos2 are conditional growth mutants necessary for eye development

We previously reported on the non-autonomous overgrowth of the *Ptc* ^*B.2.13*^, *Dark*^*82*^ mosaic eyes as compared to the control *Dark*^*82*^ mosaic eyes (Figure 1A compared to 1B, mutant tissue is pigmented) (Kagey et al., 2012). Here we add that *Cos2* ^*F.1.4*^, *Dark*^*82*^ also results in dramatically overgrown mosaic eye when compared to the control mosaic *Dark*^*82*^ (Figure 1A compared to 1C, mutant tissue pigmented). To visualize the ommatidial organization of the eye we mated mutants to an *FRT42D* chromosome with GFP (*Ey-Flp;FRT42D, ubi-GFP*).

**Figure 1:**
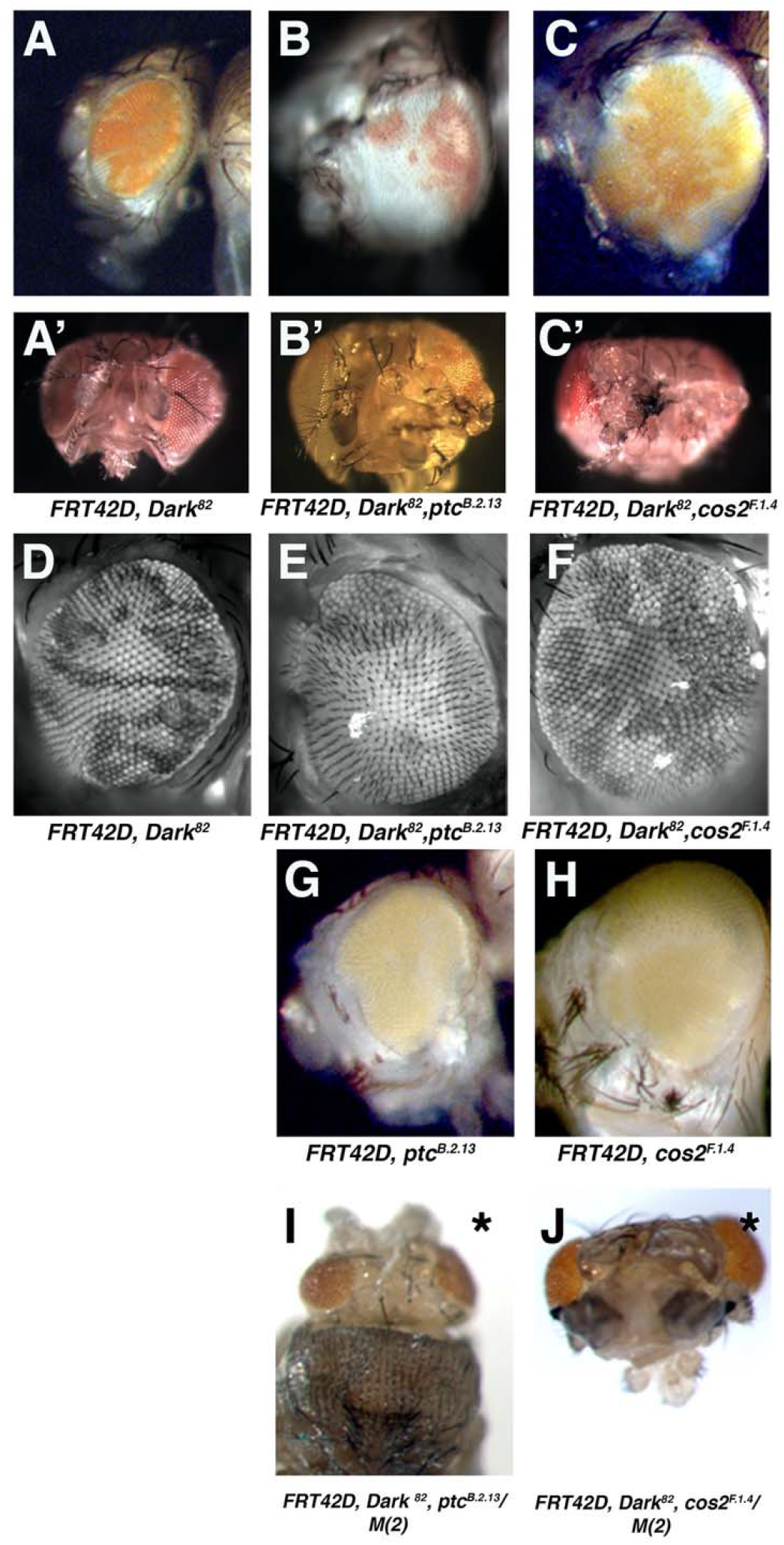
Alleles of *Patched* and *Costal-2* isolated from the same conditional Flp/FRT screen as negative regulators of cell growth and cell division in the eye. Mosaic adult eyes imaged via light microscopy by mating *Ey-Flp;FRT42D* to (A) control eye *FRT42D, Dark*^*82*^ (B) *;FRT42D, ptc*^*B.2.13*^, *Dark*^*82*^ (C) *FRT42D, Cos2*^*F.1.4*^, *Dark*^*82*^ (mutant tissue pigmented *mw+*). Dorsal images of each genotype showed in A’-C’. Fluorescent apotome images to visualize ommatidial organization by mating *Ey-Flp;FRT42D, ubi>GFP* to (D) control eye *;FRT42D, Dark*^*82*^ (E) *;FRT42D, ptc*^*B.2.13*^, *Dark*^*82*^ (F) *;FRT42D, Cos2*^*F.1.4*^, *Dark*^*82*^ (mutant tissue, darker, fluorescent negative). Light microscope images without *Dark*^*82*^ allele through mating *Ey-Flp;FRT42D, ubi-GFP* to (G) *;FRT42D, ptc*^*B.2.13*^ (H) *;FRT42D, Cos2*^*F.1.4*^ (mutant tissue is non-pigmented). Light microscope images of heads comprised entirely of mutant tissue by mating *Ey-Flp;FRT42D, M(2)* to (I) *FRT42D, Ptc*^*B.2.13*^, *Dark*^*82*^ (J) *;FRT42D, Cos2*^*F.1.4*^, *Dark*^*82*^ (mutant tissue is pigmented *mw+*). * denotes pupal lethality

Fluorescent apotome images show the *Dark*^*82*^ mosaic eye to have organized and patterned ommatidial structure in both the *Dark*^*82*^*/Dark*^*82*^ clones (darker tissue) and adjacent wild type clones (brighter GFP positive tissue) (Figure 1D). In contrast, both *Dark*^*82*^, *Ptc*^*B.2.13*^ and *Dark*^*82*^, *Cos2*^*F.1.4*^ have a disrupted ommatidial organization that can be visualized by the misalignment of ommatidia throughout the eye, the disruption of ommatidial organization can be seen in both mutant clones (darker tissue) and wild type clones (lighter tissue) suggesting that Hh pathway disruption has both autonomous and non-autonomous effects on overall eye development (Figure 1E-F,).

To investigate the degree to which the block in apoptosis contributes to the overgrowth phenotypes we reintroduced the wild type *Dark* allele through recombination, allowing homozygous cells to again go through apoptosis in mosaic eyes. Without a block in apoptosis, both *Ptc* ^*B.2.13*^ and *Cos2* ^*F.1.4*^ mosaic eyes depict a partial rescue of overgrowth, indicating that both mutant overgrowth phenotypes are, at least in part, dependent upon a block in apoptosis (Figure 1G-H, mutant tissue unpigmented). Though the eyes size is reduced when apoptosis is reintroduced, eye organizational disruption remains and can be observed in both Hh mutants. Additionally, neither *Ptc*^*B.2.13*^ or *Cos2*^*F.1.4*^ mosaic eyes had any visible unpigmented tissue (mutant tissue) cells which indicates that the majority (if not all) of the mutant cells may have been cleared by apoptosis.

To look at eyes that are entirely comprised of mutant tissue we mated the *Ptc*^*B.2.13*^ and *Cos2*^*F.1.4*^ mutants to an FRT chromosome with a homozygous lethal mutation. In both cases *Ptc*^*B.2.13*^, *Dark*^*82*^ and *Cos2*^*F.1.4*^, *Dark*^*82*^ mutants resulted in pupal lethality and a severe reduction in overall head size, highlighting the need for functional Hedgehog signaling in the developing eye and overall organismal survival (Figure 1I-J).

### Ptc and Cos2 mutations drive conditional overgrowth and pupal lethality in the mosaic wing

To investigate the generalized nature of this conditional Hh-deregulated overgrowth, we analyzed mosaic wing discs phenotypes utilizing the *UBX-Flp* driver. Previously, we found that *Ptc*^*B.2.13*^, *Dark*^*82*^ mosaic wing led to a dramatic tissue overgrowth, resulting in pupal lethality due to wing size (Kagey et al., 2012). Here we find that similar to *Ptc* ^*B.2.13*^, *Dark*^*82*^ (Figure 2B), the *Cos2*^*F.1.4*^, *Dark*^*82*^ mosaic wing discs result in substantial tissue overgrowth and complete pupal lethality (Figure 2A compared to 2C). While both the *Ptc*^*B.2.13*^, *Dark*^*82*^ and *Cos2* ^*F.1.4*^, *Dark*^*82*^ mosaic wing discs were substantially larger than the *Dark*^*82*^ control disc, we noted that the overall size of the *Ptc* ^*B.2.13*^, *Dark*^*82*^ mosaic disc was consistently larger than the *Cos2*^*F* .*1.4*^, *Dark*^*82*^ mosaic wing disc, though the difference was not statistically significant (p=0.06355497)(Figure 2D). Despite there not being a statistical difference in size, there was an observable biological difference between *Ptc*^*B.2.13*^ and *Cos2*^*F.1.4*^ mutants when the ability to undergo apoptosis was re-introduced by recombining the wild type *Dark*^*WT*^ allele. The reintroduction of apoptosis (via *Dark*^*+*^) fully rescued the *Cos2* ^*F.1.4*^ mosaic pupal lethality, while the majority of *Ptc* ^*B.2.13*^ mosaic organisms still succumbed to pupal lethality (Figure 2E). Furthermore, the resulting *Cos2* ^*F.1.4*^ mosaic wings were indistinguishable in size from control mosaic wings (Figure 2F). The rare *Ptc*^*B.2.13*^ escaper still had enlarged adults wing and depicted the ‘wings held out’ phenotype (Figure 2G). So while both *Ptc* ^*B.2.13*^ and *Cos2*^*F.1.4*^ result in dramatic wing overgrowth there are differences in the extent of tissue overgrowth based on which part of the Hh pathway is dirsrupted.

**Figure 2:**
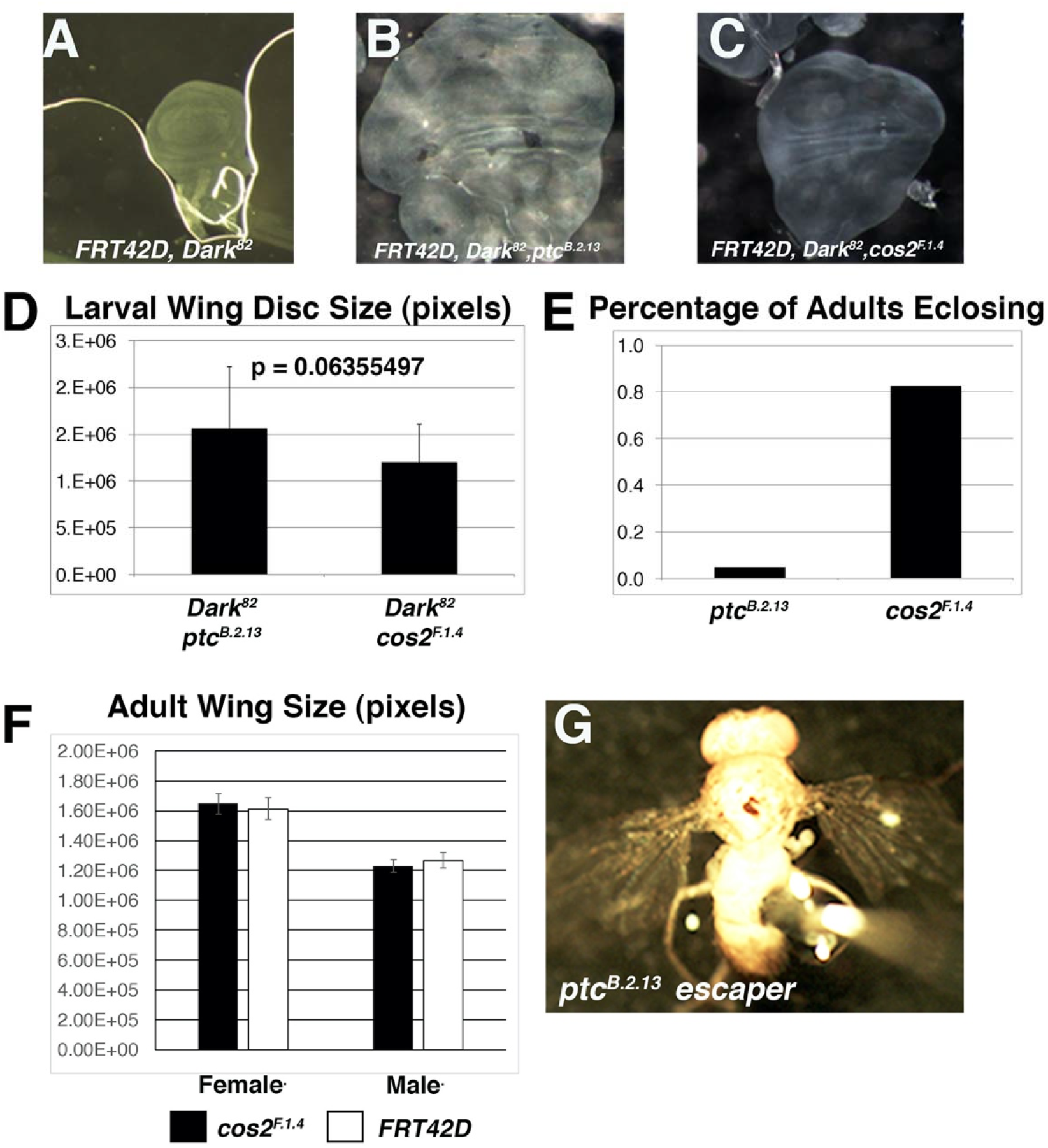
*Ptc*^***B.2.13***^ **and *Cos2***^***F.1.4***^ **exhibit conditional overgrowth phenotypes in the *Drosophila* wing that result in pupal lethality.** Third instar imaginal wing discs were visualized by light microscopy for *UBX-Flp:FRT42D* mated to (A) control eye *FRT42D, Dark*^*82*^ (B) *FRT42D, ptc*^*B.2.13*^, *Dark*^*82*^ (C) *FRT42D, Cos2*^*F.1.4*^, *Dark*^*82*^. Both *Ptc* and *Cos2* wing discs resulted in complete pupal lethality. Average size of third instar imaginal wing disc (in pixels) for (D) *FRT42D, ptc*^*B.2.13*^, *Dark*^*82*^ and *FRT42D, Cos2*^*F.1.4*^, *Dark*^*82*^ mosaic wing discs. Error bars represent standard deviation. Difference is not statistically significant. (E) Percentage of adults that eclosed for without *Dark*^*82*^ allele for *FRT42D, ptc*^*B.2.13*^ and *FRT42D, Cos2*^*F.1.4*^ (n=100 for each genotype). (F) Comparison of adult wing size (in pixels) of control (*FRT42D*) and (*FRT42D, Cos2*^*F.1.4*^) wings, 10 wings per genotype and sex. (G) Image of *FRT42D, ptcB.2.13* escaper, depicting wings held out phenotype.

### Patched and Costal-2 exhibit differences in the autonomy of overgrowth

While both *Ptc*^*B.2.13*^ and *Cos2*^*F.1*.4^ are conditional regulators of cell growth and tissue size, we observed differences in the ratio of mutant (pigmented) to wild type (unpigmented) tissue (Figure1B, 1C). The *Dark*^*82*^, *Ptc* ^*B.2.13*^ mosaic eye is comprised of mostly wild-type tissue, suggesting a non-autonomous overgrowth (Figure1B, 1E and (Kagey et al., 2012)), while the *Dark*^*82*^, *Cos2* ^*F.1.4*^ mosaic eye depicts more gross overgrowth (both autonomous and non-autonomous tissue overgrowth Figure 1C, 1F). To measure the extent of this difference, we utilized imaginal eye and wing discs to determine the percentage each disc is comprised of homozygous mutant tissue in *Ptc*^*B.2.13*^ and *Cos2*^*F.1.4*^ mutants. Using imaginal discs allowed for quantification of 2-D tissue instead of 3-D eye ratios. In these imaginal discs we observe a statistically significant difference (p=0.01827498) in which the *Dark*^*82*^, *Ptc* ^*B.2.13*^ eye discs are comprised of (8.1%) mutant tissue (non-GFP positive), while the *Dark*^*82*^, *Cos2* ^*F.1.4*^ eye discs are comprised of (17.2%) mutant tissue (non-GFP positive)(Figure 3A-C). This difference in autonomous overgrowth despite both genotypes resulted in premature differentiation (and mitotic arrest) in the imaginal eye discs (observed by Elav expression in mutant clones before the morphogenetic furrow, Supplemental Figure 2). Similarly, in the wing disc we find that 12.6% of *Dark*^*82*^, *Ptc* ^*B.2.13*^ imaginal wings discs are comprised of mutant tissue while 25.9% of *Dark*^*82*^, *Cos2* ^*F.1.4*^ imaginal wing discs are comprised of mutant tissue, p=0.00605447 (Figure 3D-F). As a comparison, we have previously reported the Dark82 mosaic wing disc to be ∼ 38% *Dark*^*82*^*/Dark*^*82*^ mutant cells. The difference in tissue autonomy of *Ptc* and *Cos2* in the eye and wing overgrowth autonomy suggest that these are pathway disruption differences and not a phenomenon of tissue specific developmental signaling. To understand how mutations in the same pathway could lead to different varieties of tissue overgrowth, we utilized imaginal disc staining to observe molecular alterations that accompany these mutations.

**Figure 3:**
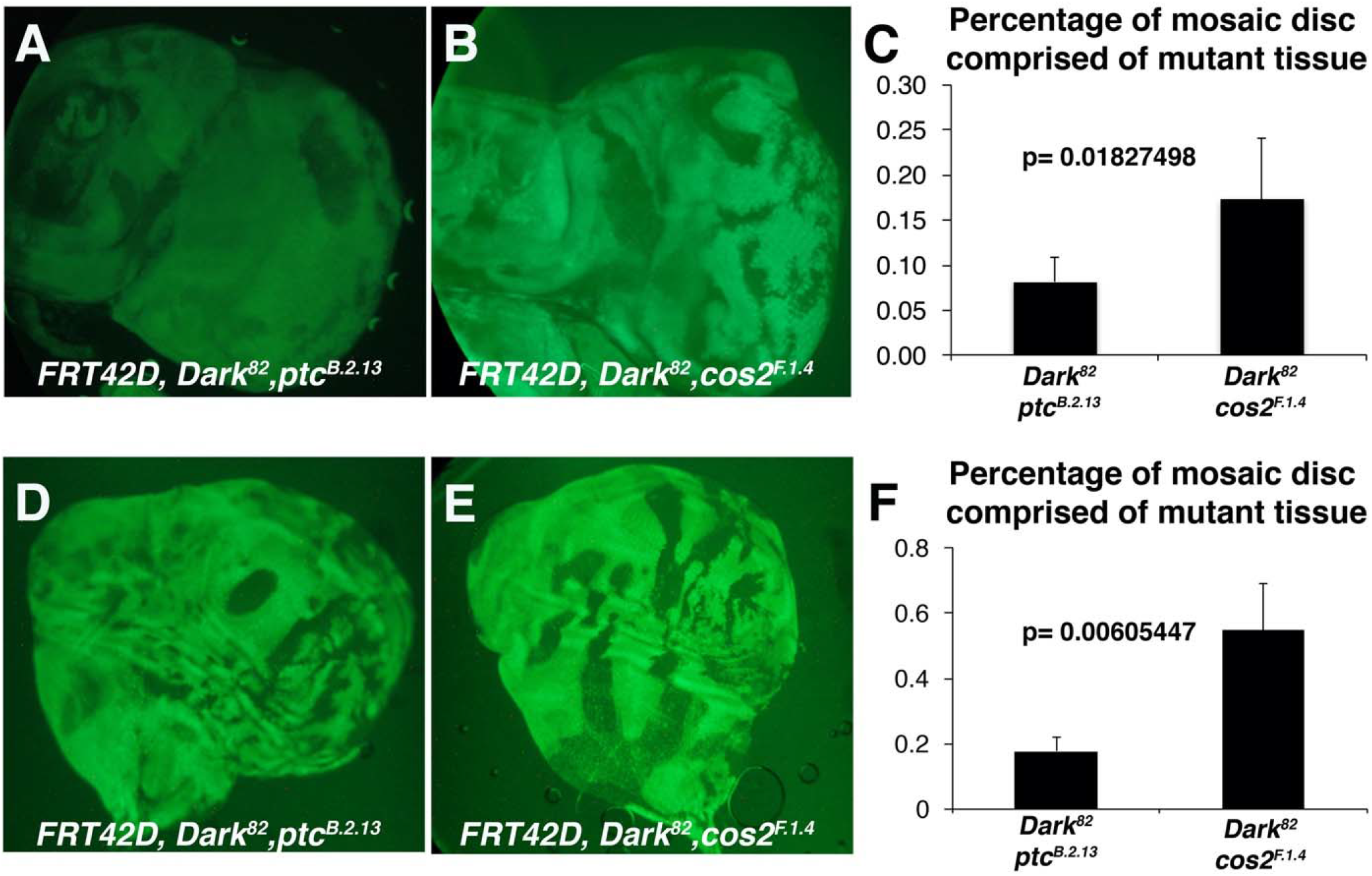
*Ptc*^***B.2.13***^ **and *Cos2***^***F.1.4***^ **exhibit distinct types of autonomous overgrowth in the eye and wing.** Third instar imaginal eye discs were visualized by fluorescent microscopy for crosses of *Ey-Flp:FRT42D, ubi-GFP* mated to (A) *FRT42D, ptc*^*B.2.13*^, *Dark*^*82*^ (B) *FRT42D, Cos2*^*F.1.4*^, *Dark*^*82*^ (homozygous mutant tissue is GFP negative). (C) Percentage of mosaic disc comprised of mutant tissue for *FRT42D, ptc*^*B.2.13*^, *Dark*^*82*^ and *FRT42D, Cos2*^*F.1.4*^, *Dark*^*82*^ measured by pixels. >10 imaginal discs quantified for each genotype. Error bars represent standard deviation and p value is from paired T-test. Third instar imaginal wing discs were visualized by light microscopy for crosses of *UBX-Flp:FRT42D, ubi-GFP* mated to (D) *FRT42D, ptc*^*B.2.13*^, *Dark*^*82*^ (E) *FRT42D, Cos2*^*F.1.4*^, *Dark*^*82*^ (homozygous mutant tissue is GFP negative). (F) Percentage of mosaic disc comprised of mutant tissue for *FRT42D, ptc*^*B.2.13*^, *Dark*^*82*^ and *FRT42D, Cos2*^*F.1.4*^, *Dark*^*82*^ measured by pixels. >10 imaginal discs quantified for each genotype. Error bars represent standard deviation and p value shown from paired T-test.

### Ptc and Cos2 autonomously de-regulate Hedgehog pathway signaling

Both Ptc and Cos2 are upstream negative regulators of the canonical Hedgehog signaling pathway and function to sequester the transcription factor Cubitus interruptus (Ci) in the cytoplasm (Aza-Blanc et al., 1997). Previously, we established that *Ptc*^*B.2.13*^ mutant clones result in an autonomous upregulation of Dpp, Patched, and Ci (Figure 4B, (Kagey et al., 2012)). *Cos2* mutations have also been shown to autonomously upregulate downstream Hh targets (Christiansen et al., 2012; Christiansen et al., 2013; Zadorozny et al., 2015). Here we support these findings through the observation of an upregulation of cleaved Ci in *Dark*^*82*^, *Ptc* ^*B.2.13*^ and *Dark*^*82*^, *Cos2* ^*F.1.4*^ imaginal wing disc clones (Figure 3B-C). Conversely, *Dark*^*82*^ mosaic wings demonstrate a wild type expression pattern of Ci suggesting that the Ci overexpression is not resultant from a block in apoptosis (Figure 3A). The deregulation of Ci in *Ptc* and *Cos2* mutants occurs exclusively in clones on the anterior side of the wing disc correspondent with the expression domain of Ci (all wing discs are oriented with anterior to the left). *Dark*^*82*^, *Cos2* ^*F.1.4*^ also phenocopies the *Dark*^*82*^, *Ptc* ^*B.2.13*^ autonomous increase of Patched expression in the anterior domain of the wing disc (Ptc is also a Ci target gene) (Supplemental Figure 3). Overall, this suggests that mutations in *Ptc* or *Cos2* mutations disrupt the Hh pathway and lead to an autonomous increase of Hh signaling.

**Figure 4:**
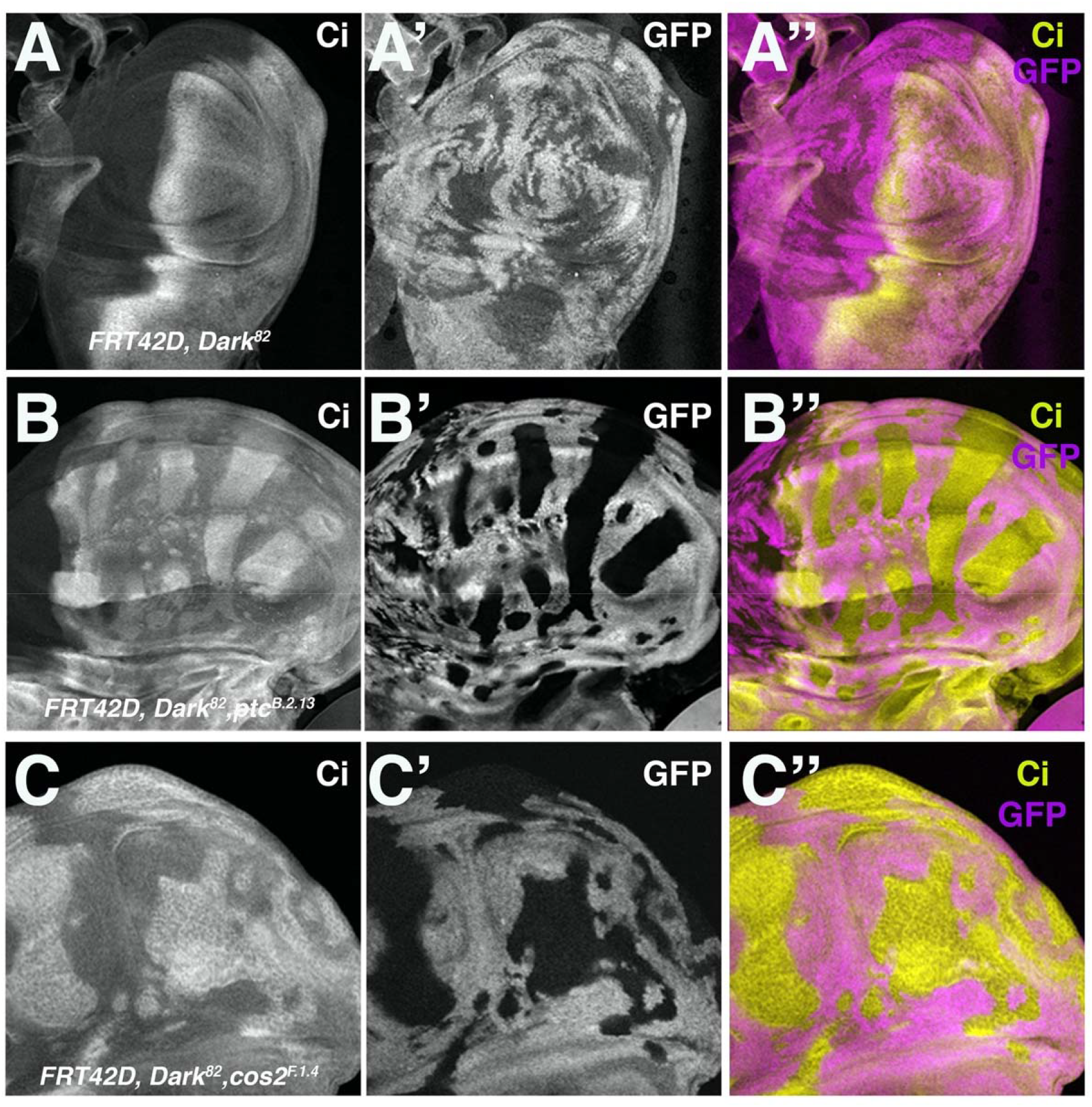
*Ptc*^***B.2.13***^ **and *Cos2***^***F.1.4***^ **autonomously up-regulated Ci within mutant clones in the anterior compartment of imaginal wing discs**. Ci levels visualized in third instar imaginal wing discs through staining and fluorescent microscopy for crosses of *UBX-Flp:FRT42D, ubi-GFP* mated to (A) control *FRT42D, Dark*^*82*^ (B) *FRT42D, ptc*^*B.2.13*^, *Dark*^*82*^ (C) *FRT42D, Cos2*^*F.1.4*^, *Dark*^*82*^ (mutant tissue is GFP negative). Anterior portion of wing discs are oriented to the right.

### Ptc and Cos2 mutant clones result in a non-autonomous up-regulation of DIAP1

Previously, our lab and others, have established a link between Hh deregulation and non-autonomous DIAP1 up-regulation in both *Ptc* and *Cos2* mosaic tissue (Christiansen et al., 2012; Christiansen et al., 2013; Kagey et al., 2012). To ensure this non-autonomous up regulation of DIAP1 is consistent across *Cos2* alleles, we measured the levels of DIAP1 expression in mosaic imaginal wing discs for *Cos2*^*F.1.4*^. As conveyed previously, *Dark*^*82*^, *Ptc* ^*B.2.13*^ leads to the non-autonomous upreguation of DIAP1 (Figure 5A), which can be visualized as the halo of DIAP1 expression just outside of the *Ptc*^*B.2.13*^ clonal borders. The *Dark*^*82*^, *Cos2* ^*F.1.4*^ mutant also leads to the non-autonomous increase in DIAP1 expression (Figure 5B) visualized as halos of DIAP1 up-regulation immediately outside the *Cos2*^*F.1.4*^ clones. To more easily visualize the non-autonomous nature of the DIAP1 overexpression we utilized Ci to mark the mutant clones which provides an easier observation of expression at the clonal boundaries. Further, this up-regulation of DIAP1 is confined to the anterior compartment suggesting it is dependent on the Ci overexpression autonomously to facilitate the non-autonomous DIAP1 up-regulation (Figure 5).

**Figure 5:**
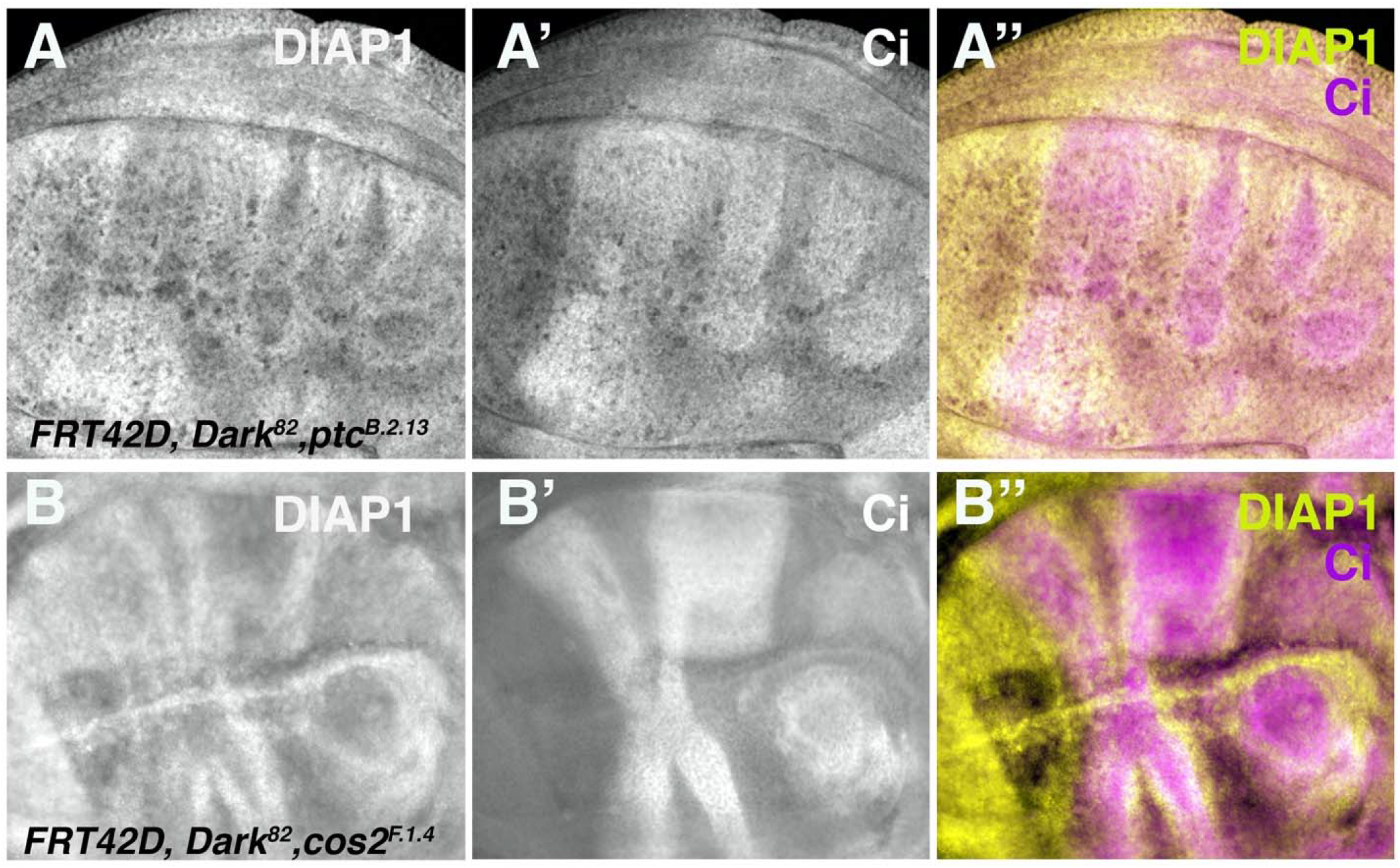
*Ptc*^***B.2.13***^ **and *Cos2***^***F.1.4***^ **non-autonomously up-regulated DIAP1 just outside of mutant clone boundaries in the anterior compartment of imaginal wing discs**. DIAP1 levels visualized in third instar imaginal wing discs through staining and fluorescent microscopy for crosses of *UBX-Flp:FRT42D, ubi-GFP* mated to (A) *FRT42D, ptc*^*B.2.13*^, *Dark*^*82*^ (B) *FRT42D, Cos2*^*F.1.4*^, *Dark*^*82*^ (mutant tissue are marked by Ci up-regulated tissue). Anterior portion of wing discs are oriented to the right.

### Ptc and Cos2 mutations have different patterns of de-regulated pMad1/5

Previously, we found that the non-autonomous increase in DIAP1 and non-autonomous tissue proliferation of the *Dark*^*82*^, *Ptc* ^*B.2.13*^ mutants were shown to be dependent on the concurrent non-autonomous up regulation of pMad1/5 and the interaction of pMad and Yorkie to drive non-autonomous survival and proliferation (Kagey et al., 2012). Here we investigated if *Cos2*^*F.1.4*^ mutant clones also de-regulate pMad in a similar manner. The non-autonomous activation of pMad1/5 in *Dark*^*82*^, *Ptc* ^*B.2.13*^ mosaic wing discs, is visualized by the demonstrable increase in pMad expression just outside of the mutant clone (mutant clones again marked by Ci expression to visualize pMad expression at the clonal boundaries) (Figure 6A-A’’). This non-autonomous increase is also seen via line scan, where the peaks of pMad expression are just outside of the mutant clonal boundaries and dissipate further away from the mutant clone (Figure 6B, clones marked by arrows).

**Figure 6:**
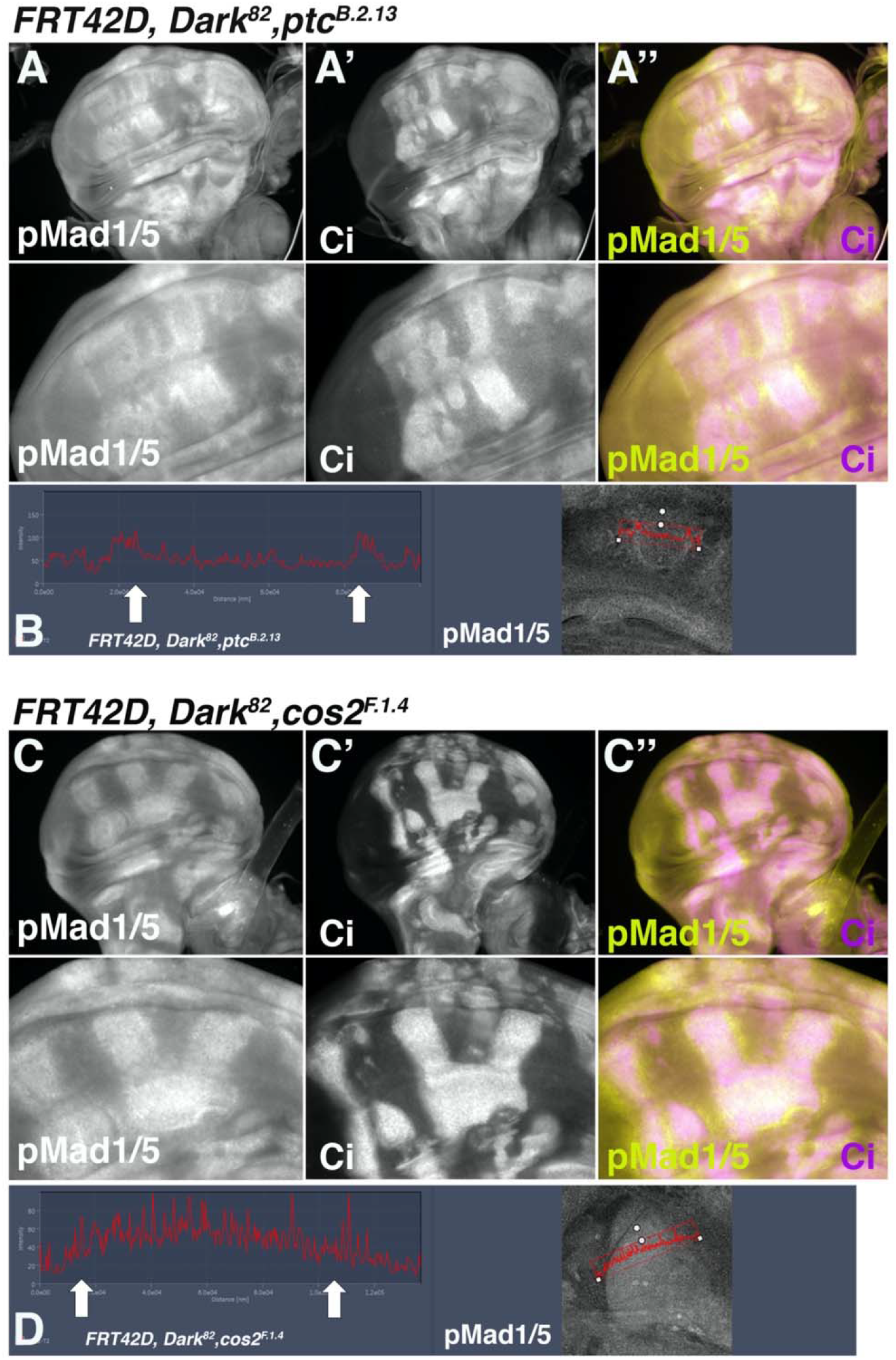
*Ptc*^***B.2.13***^ **and *Cos2***^***F.1.4***^ **differentially up-regulate pMad1/5 inside of mutant clones**. pMad 1/5 levels visualized in third instar imaginal wing discs through staining and fluorescent microscopy for crosses of *UBX-Flp:FRT42D, ubi-GFP* mated to (A-A’’) *FRT42D, Ptc*^*B.2.13*^, *Dark*^*82*^ (mutant tissue are marked by Ci up-regulated tissue) (B) quantification of pMad1/5 levels across the mutant clonal boundaries in *Ptc*^*B.2.13*^ mutant clones. Arrows denote clonal boundaries. (C-C’’) *FRT42D, Cos2*^*F.1.4*^, *Dark*^*82*^ (mutant tissue are marked by Ci up-regulated tissue) (D) quantification of pMad1/5 levels across the mutant clonal boundaries in *Ptc*^*B.2.13*^ mutant clones. Arrows denote clonal boundaries. Anterior portion of wing discs are oriented to the right.

In *Dark*^*82*^, *Cos2* ^*F.1.4*^ clones we also see a de-regulation and overexpression pMad expression. However, in contrast to the purely non-autonomous expression seen in the *Ptc*^*B.2.13*^ clones, in *Cos2*^*F.1.4*^ the up-regulation of pMad is both autonomous (within the clone) and non-autonomous (adjacent to the clone) (Figure 6C’-C’’, mutant clones marked by Ci to visualize boundary expression). We again used a line scan to visualize the pMad expression intensity. The high levels of pMad expression came from within the mutant clone and then remained high outside of the clonal boundary (Figure 6D, clonal boundaries arrows). This indicates that while both *Ptc* and *Cos2* mutant tissue lead to a non-autonomous increase of pMad, only *Cos2* has an autonomous upregulation of pMad expression. The regions which pMad overexpression is observed correlate to the nature of tissue overgrowth, non-autonomous, for *Ptc*^*B.2.13*^ and both autonomous and non-autonomous for *Cos2*^*F.1.4*^.

## Discussion

### Overview

We designed a genetic screen to identify negative regulators of cell growth and cell division that were conditional on a need for a block in apoptosis. From this screen, we isolated and mapped a novel allele of *Cos2, Cos2*^*F.1.4*^, as a conditional growth regulator from an EMS Flp/FRT mosaic eye screen. This finding builds upon previous data where we identified another member of the hedgehog signaling pathway, *Ptc*^*B.2.13*^, in the same screen, providing confirmation that disruption in the hedgehog pathway can lead to conditional tissue overgrowth in *Drosophila* (Kagey et al., 2012). While both *Ptc* and *Cos2* mutants result in dramatic eye and wing overgrowth due to the autonomous disruption of the Hh pathway, we observed that the nature of the overgrowth differs between *Ptc* and *Cos2* mutants. *Ptc* mosaic tissue displays a distinct non-autonomous overgrowth whereas *Cos2* mosaic tissue results in both autonomous and non-autonomous overgrowth. These differences were consistent in both the eye and wing imaginal discs. Our data indicate that this difference may be due to the differential pattern of pMad deregulation. The up-regulation of pMad has previously been demonstrated to drive tissue overgrowth (Kagey et al., 2012; Oh and Irvine, 2011), and the pattern of pMad deregulation in *Ptc* and *Cos2* mosaic tissue directly matches the patterns of overgrowth.

### Association of cell survival and Hedgehog signaling

Our results suggest that a functional Hh signaling pathway is necessary for cell survival in imaginal disc development. Mutations to the *Ptc* and *Cos2* members of the hedgehog pathway led to cell death. When apoptosis was blocked, *Ptc* and *Cos2* mutants depicted dramatic overgrowth, which was rescued in both the eye and wing when the wild type *Dark* allele was reintroduced, thus reinstating cell death.

This is further supported by the findings of the Bergman lab that identified multiple members of the Hh pathway in the *GMR-Hid Flp/FRT* screen (Christiansen et al., 2012; Christiansen et al., 2013; Fan and Bergmann, 2008). This study also observed an autonomous increase in cell death when the Hh pathway is disrupted. The finding that this cell death is accompanied by a concurrent non-autonomous survival signal may be due to the compensatory proliferation resulting from the loss of Hh-dependent cell death (Worley et al., 2012). This concurrent non-autonomous DIAP1 and proliferation increase could be a mechanism of tissue regeneration from which dying cells signal to their neighbors to proliferate. In our scenario, we have removed the ability of the Hh deficient cells to die through the canonical apoptotic pathway, therefore creating a constitutive non-autonomous survival and growth signaling leading to eye and wing overgrowth.

### Ptc and Cos2 exhibit different patterns of pMad deregulation

Both *Ptc* and *Cos2* mutations lead to an autonomous increase in hedgehog signaling (including Ci, and Ptc); however, there may be a difference in which type of cells can interpret the increased Hh target Dpp. Given that Dpp is a morphogen, both autonomous and non-autonomous cells would receive the Dpp signal via the receptors Thick Veins and Punt (Hamaratoglu et al., 2014). However, as we previously demonstrated, the *Ptc*^*B.2.13*^ mutant clones also autonomously upregulated the inhibitory smad, Daughters Against Decapentaplegic (Dad) (Goldstein et al., 2011; Kagey et al., 2012). We hypothesize that a lack of Dad upregulation in the *Cos2*^*F.1.4*^ mutant clones would result in an autonomous increase proliferation due to the Dpp mediated pMad activation (as seen in Figure 6). Alternatively, it is possible that a differential downregulation of Tkv downstream of the Hh signaling could be leading to the differential activation of pMad between *Ptc* and *Cos2* mutant clones (Tanimoto et al., 2000). In either scenario, the difference in autonomous pMad upregulation between *Ptc* and *Cos2* mosaic tissue highlights the importance of understanding how disrupting different points of conserved pathways impacts tissue patterning and growth.

### Potential implications for understanding Hh disrupted human tumors

A detailed understanding of the precise genetic mechanisms and pathways leading to de-regulated growth in Hh mutations will be important in developing diagnostic and treatment tools to address Hh de-regulated tumors in humas. Hedgehog pathway mutations are associated with various types of human cancers, including medulloblastoma; basal cell carcinoma; glioma; and breast, colorectal, pancreatic, and prostate cancer (Jiang and Hui, 2008; Skoda et al., 2018). The loss of *Ptc* also increases the occurrence of these types of tumors in mice, providing additional models for study (Wu et al., 2011). There is evidence of both Ptc and Kif7 (the human homolog of Cos2) being down-regulated in different human cancers, including basal cell carcinoma and ovarian cancer (Yao et al., 2019). Fully understanding the phenotypic differences that arise from disruptions at different points in the Hh pathway may ultimately help with personalized medicine and treatment decisions for patients with Hh-deregulated tumors.

## Acknowledgements

We thank A. Bergmann, T. Cook, and K. Moberg for *Drosophila* stocks and reagents. We thank T. Cook and E. Kagey for helpful comments on earlier drafts of this manuscript. Antibodies were obtained from the Developmental Studies Hybridoma Bank, created by the NICHD of the NIH and maintained at The University of Iowa, Department of Biology, Iowa City, IA 52242. Stocks obtained from the Bloomington Drosophila Stock Center (NIH P40OD018537) were used in this study.

---

**Supplemental Figure 1:**
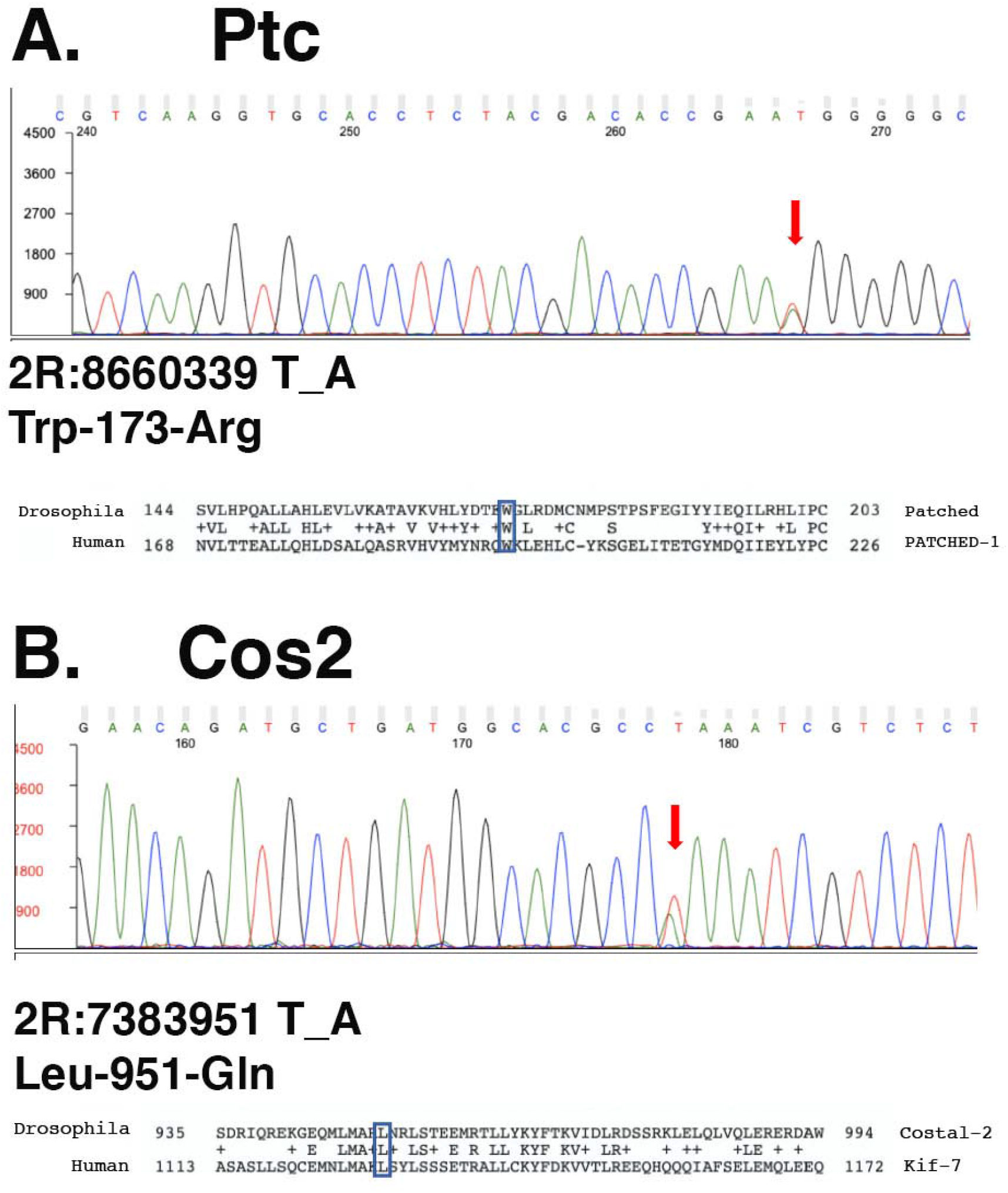
*Ptc*^***B.2.13***^ **and *Cos2***^***F.1.4***^ **have missense mutations in conserved amino acids**. *Ptc* and *Cos2* heterozygous animals were sequenced via Sanger sequencing. (A) *Ptc*^*B.2.13*^ mutation resides at 2R:8660339 resulting in a Trp-Arg mutation that is in a residue conserved between *Patched* and *PATCHED1*. (B). *Cos2*^*F.1.4*^ mutation resides at 2R: 7383951 resulting in a Leu-Gln mutation in a conserved residue between *Costal2* and *Kif7*.

**Supplemental Figure 2:**
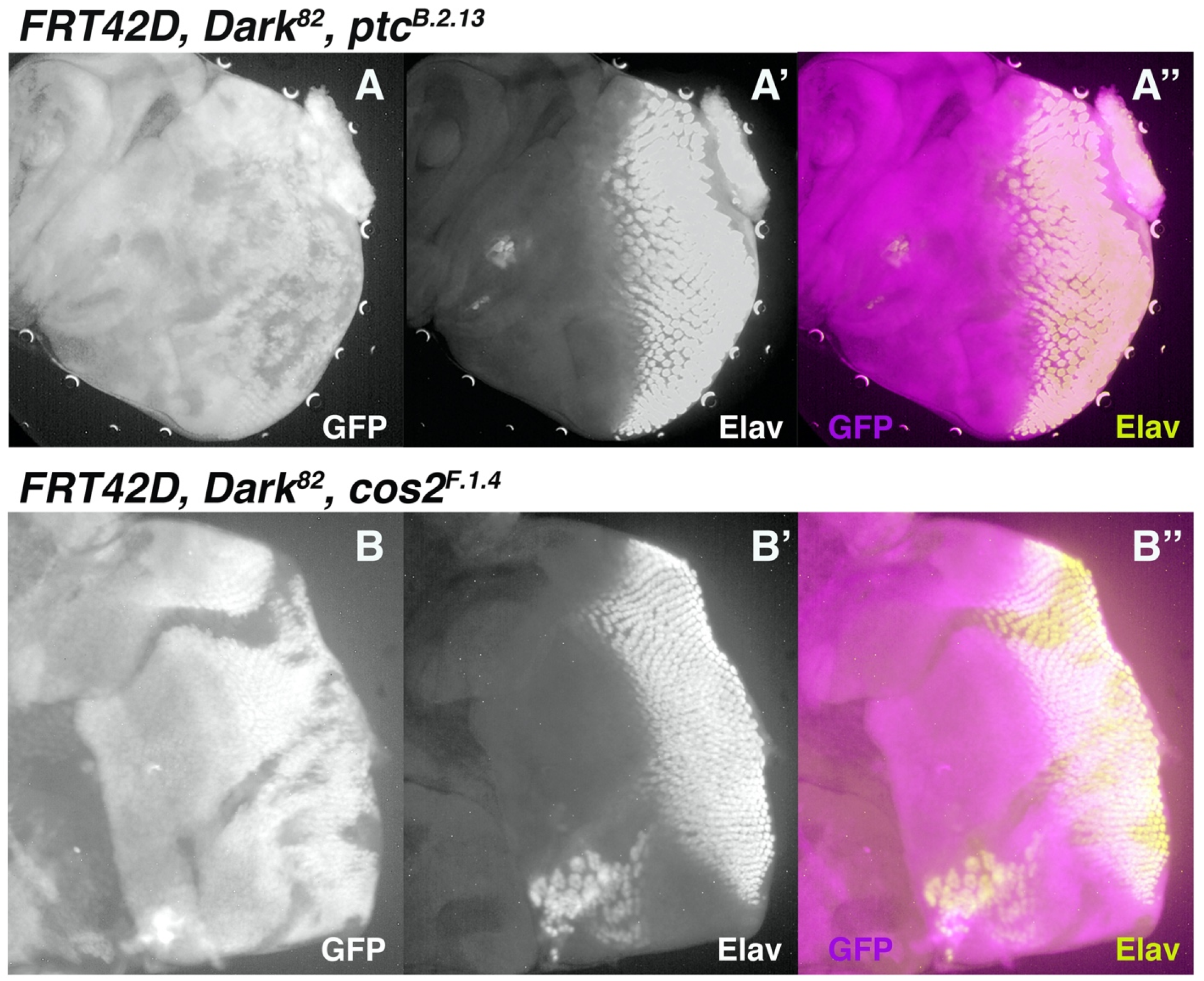
*Ptc*^***B.2.13***^ **and *Cos2***^***F.1.4***^ **clones result in premature Elav expression and eye differentiation.** Elav levels visualized in third instar imaginal eye discs through staining and fluorescent microscopy for crosses of *Ey-Flp:FRT42D, ubi-GFP* mated to (A-A”“) *FRT42D, Ptc*^*B.2.13*^, *Dark*^*82*^ and (B-B”) *FRT42D, Cos2*^*F.1.4*^, *Dark*^*82*^ (mutant tissue is GFP negative). In both mutants ectopic Elav expression can be seen in mutant clones prior to the wave of differentiation seen in the rest of the eye.

**Supplemental Figure 3:**
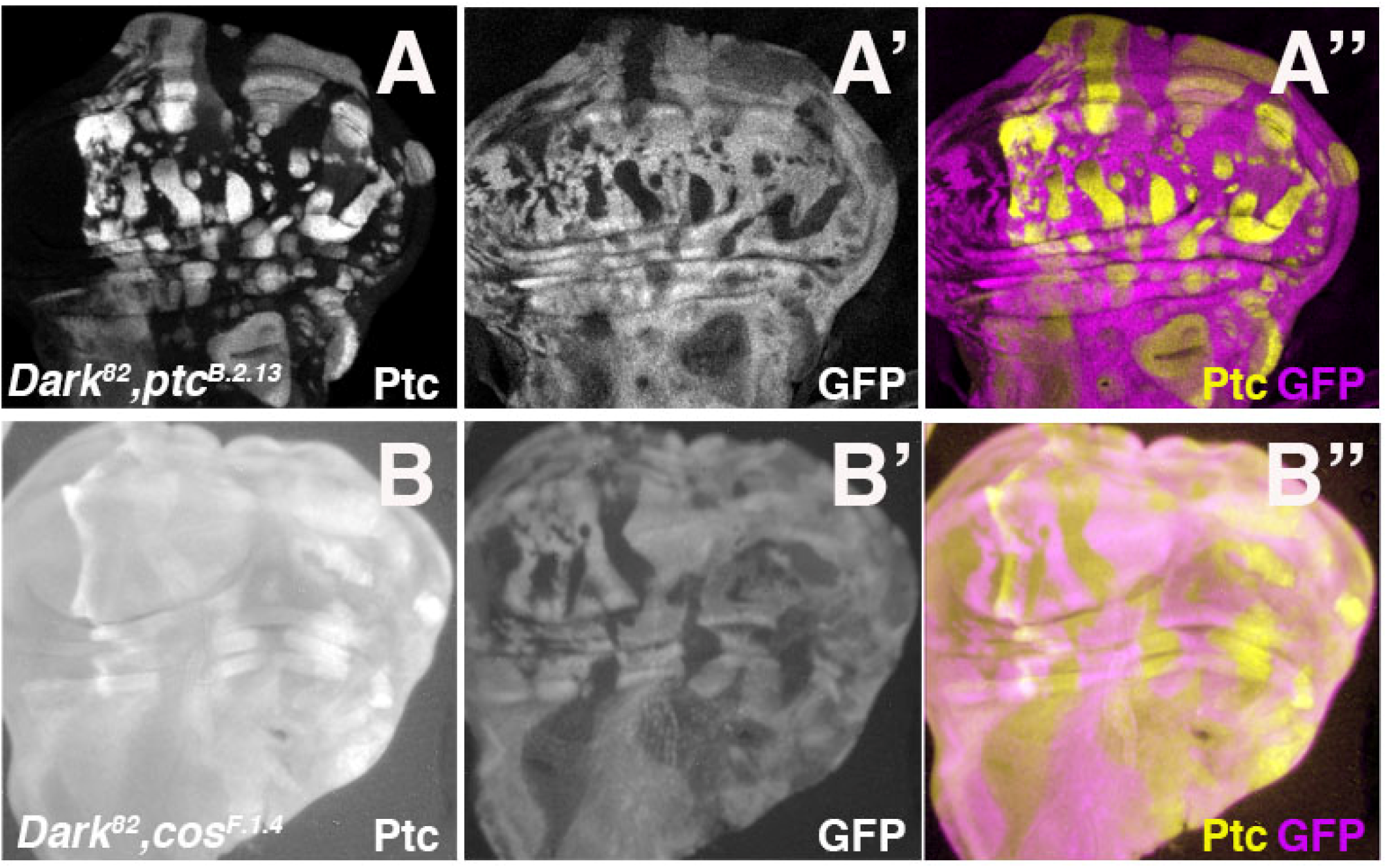
*Ptc*^***B.2.13***^ **and *Cos2***^***F.1.4***^ **autonomously up-regulated Ptc within mutant clones in the anterior compartment of imaginal wing discs**. Ptc levels visualized in third instar imaginal wing discs through staining and fluorescent microscopy for crosses of *UBX-Flp:FRT42D, ubi>GFP* mated (A-A’’) *FRT42D, Ptc*^*B.2.13*^, *Dark*^*82*^ and (B-B’’) *FRT42D, Cos2*^*F.1.4*^, *Dark*^*82*^ (mutant tissue is GFP negative). Anterior portion of wing discs are oriented to the right.

